# Impairment of methylglyoxal detoxification systems causes mitochondrial dysfunction and schizophrenia-like behavioral deficits

**DOI:** 10.1101/2020.07.08.192906

**Authors:** Kazuya Toriumi, Stefano Berto, Shin Koike, Noriyoshi Usui, Takashi Dan, Kazuhiro Suzuki, Mitsuhiro Miyashita, Yasue Horiuchi, Akane Yoshikawa, Yuki Sugaya, Takaki Watanabe, Mai Asakura, Masanobu Kano, Yuki Ogasawara, Toshio Miyata, Masanari Itokawa, Genevieve Konopka, Makoto Arai

## Abstract

Methylglyoxal (MG) is a cytotoxic *α*-dicarbonyl byproduct of glycolysis. Our bodies have several bio-defense systems to detoxify MG, including an enzymatic system by glyoxalase (GLO) 1 and a scavenge system by vitamin B6 (VB6). We know a population of patients with schizophrenia impaired MG detoxification systems. However, the molecular mechanism connecting them remains poorly understood. We created a novel mouse model for MG detoxification deficits by feeding *Glo1* knockout mice VB6-lacking diets (KO/VB6(-)) and evaluated the effects of impaired MG detoxification systems on brain function. KO/VB6(-) mice accumulated MG in the prefrontal cortex (PFC), hippocampus, and striatum, and displayed schizophrenia-like behavioral deficits. Furthermore, we found aberrant gene expression related to mitochondria function in the PFC of the KO/VB6(-) mice. We demonstrated respiratory deficits in mitochondria isolated from the PFC of KO/VB6(-) mice. These findings suggest that MG detoxification deficits might cause schizophrenia-like behavioral deficits via mitochondrial dysfunction in the PFC.

## Introduction

Methylglyoxal (MG) is a highly reactive *α*-ketoaldehyde formed endogenously as a byproduct of the glycolytic pathway, by the degradation of triphosphates or by nonenzymatic fragmentation of sugar^1^. MG accumulates under conditions of hyperglycemia, impaired glucose metabolism, or oxidative stress. An excess of MG formation causes mitochondrial impairment and reactive oxygen species (ROS) production that further increases oxidative stress. It also leads to the formation of advanced glycation end products (AGEs) due to MG reacting with proteins, DNA, and other biomolecules^2, 3^. This implies accumulating MG causes damage in various tissues and organs^4^, resulting in aging and diabetic complications, such as neuropathy, retinopathy, and ischemic heart disease. Also, AGEs formed by MG induce aberrant inflammation via binding to receptor for AGEs (RAGE), which play a role in chronic inflammation and Alzheimer’s disease^5, 6^.

To remove the toxic MG, various detoxification systems work together *in vivo*. The glyoxalase system comprised of two enzymes, glyoxalase (GLO) 1 and GLO2 is an enzymatic pathway that catalyzes the glutathione-dependent detoxification of MG. GLO1 catalyzes the conversion of the hemithioacetal formed by the nonenzymatic reaction of glutathione with MG, to S-D-lactoylglutathione, which is then converted to D-lactate by GLO2^1, 7^. GLO1 and GLO2 are ubiquitously expressed in various tissues—including the brain—and provide an effective defense against the accumulations of MG. Another MG detoxification system is a carbonyl scavenge system carried out by vitamin B6 (VB6). VB6 can directly scavenge MG and trap ROS^8, 9^. Additionally, VB6 can inhibit AGE formation by blocking oxidative degradation of the Amadori intermediate of the Maillard reaction^10, 11^.

Schizophrenia is a heterogeneous psychiatric disorder characterized by positive symptoms, such as hallucinations and delusions, negative symptoms, such as anhedonia and flat affect, and cognitive impairment. We have reported that several patients with schizophrenia have a novel heterozygous frameshift and a single nucleotide variation (SNV) in *GLO1* that results in reductions of enzymatic activity. Furthermore, we have reported that VB6 (pyridoxal) levels in peripheral blood of patients with schizophrenia are significantly lower than that of healthy controls^12–14^. More than 35% of patients with schizophrenia have low levels of VB6 (clinically defined as male: < 6 ng/ml, female: < 4 ng/ml). Other groups have replicated these results^15, 16^.

VB6 levels are inversely proportional to the severity score on the Positive and Negative Syndrome Scale (PANSS)^14^. Also, a recent umbrella review has shown the decreased VB6 in patients with schizophrenia as the most convincing evidence in peripheral biomarkers for major mental disorders^17^. Considering their protective functions in detoxifying MG, the *GLO1* malfunction and the VB6 deficiency observed in patients with schizophrenia would induce the accumulation of MG. This suggests that MG accumulation may contribute to the pathophysiology of schizophrenia. Moreover, we recently reported that high-dose VB6 treatment was effective in alleviating psychotic symptoms (especially PANSS negative and general subscales), in a subset of patients with schizophrenia, coincident with decreased AGEs levels in the peripheral blood^18^. Despite this evidence, the effects of MG detoxification deficits on the pathophysiology of schizophrenia *in vivo* remain unclear.

In the present study, we generated *Glo1* KO mice to uncover the influence of MG detoxification deficits on behavioral performances and neurochemical function. We found that *Glo1* KO mice fed with VB6-lacking (VB6(-)) diets exhibited an accumulation of MG within the brain and sensorimotor deficit in a prepulse inhibition (PPI) test, although the *Glo1* gene deletion or VB6 deficiency alone did not cause the characteristic changes. Furthermore, we found aberrant gene expression involved in mitochondria function in the prefrontal cortex (PFC) of the *Glo1* KO mice with VB6 deficiency and we demonstrated respiratory impairment in mitochondria within the PFC of *Glo1* KO mice.

## Results

### Generation of Glo1 knockout mouse

We generated *Glo1* KO mice by gene trapping (**Fig. 1A**) and confirmed the insertion of the gene-trap cassette into the *Glo1* gene by PCR (**Fig. 1B**). We confirmed *Glo1* mRNA decreased to approximately 50 % and 0 % in the *Glo1* HE and KO mice, respectively, without any changes in gene expression of other enzymes for MG detoxification such as glyoxalase 2 (*Glo2*) and aldo-keto reductase (*Akr*) (**Fig. 1C**). We also determined the loss of the GLO1 protein in various organs and brain regions of *Glo1* KO mice (**Fig. 1D and 1E**) and loss of GLO1 enzymatic activity in the whole brain of *Glo1* KO mice (**Fig. 1F**). Furthermore, the body weight in *Glo1* KO mice was significantly decreased in both sexes compared with WT mice (**Fig. 1G**).

**Figure 1:**
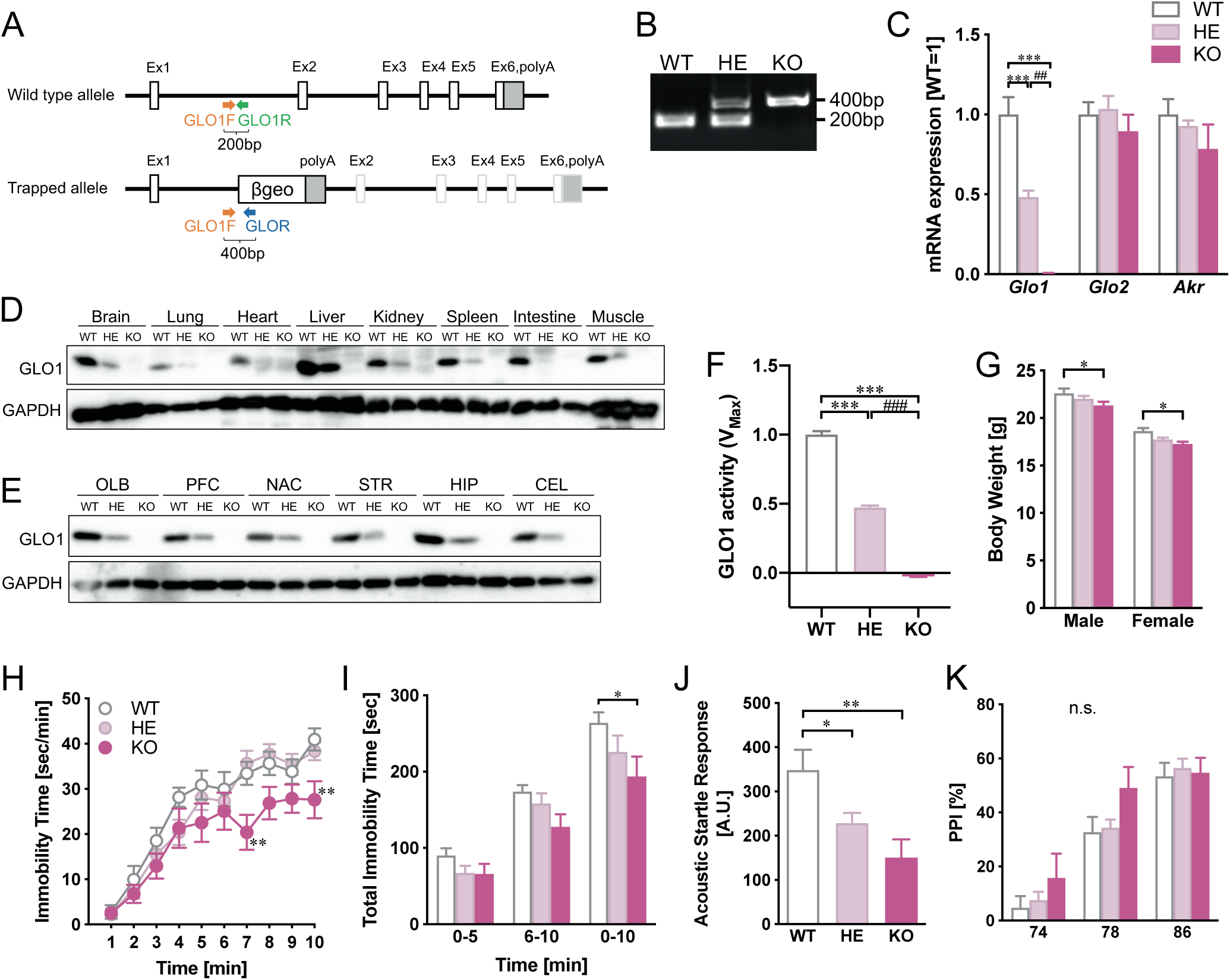
Generation of Glo1 knockout mice. (A) An overview of *Glo1* gene trapping and (B) PCR for genotyping are shown. (C) mRNA expression in the whole brain was quantified by qPCR (n = 4). Two-way ANOVA: *F*Interaction(4,18) = 13.7, *p <* 0.0001; *F*Gene(2,18) = 46.4, *p* < 0.0001; *F*Genotype(2, 9) = 9.88, *p* < 0.01. (D) In the various organs (D) and each brain regions (E), protein level of GLO1 is shown by western blotting using an anti-GLO1 antibody (Clone 6F10, Sigma) and anti-GAPDH (6C5, Santa cruz). (F) GLO1 activity was measured in whole brain (n = 3). One-way ANOVA: *F*(2,6) = 880, *p <* 0.0001. (G) Body weight was measured at 8 weeks (n = 7-19). Two-way ANOVA: *F*Interaction(2,64) = 0.168, *p >* 0.05; *F*Sex(1,64) = 250, *p* < 0.0001; *F*Genotype(2, 64) = 7.26, *p* < 0.01. In the forced swimming test, (H) the time course and (I) total scores of the immobility time are shown. Two-way ANOVA: (H) *F*Interaction(18, 396) = 1.89; *p* < 0.05, *F*Time(9, 396) = 75.9; *p* < 0.0001, *F*Genotype(2, 44) = 3.35; *p* < 0.05, (I) *F*Interaction(4, 92) = 2.08; *p* > 0.05, *F*Time(2, 92) = 248; *p* < 0.0001, *F*Genotype(2, 46) = 2.16; *p* = 0.13. In the prepulse inhibition test, (J) acoustic startle response without prepulse [One-way ANOVA: *F*(2, 41) = 6.91, *p <* 0.01.] and (K) prepulse inhibition [Two-way ANOVA: *F*Interaction(4, 84) = 2.41; *p* > 0.05, *F*Prepulse(2, 84) = 183; *p* < 0.0001, *F*Genotype(2, 42) = 0.35; *p* > 0.05.] are shown. \**p* < 0.05, \*\**p* < 0.01, \*\*\**p* < 0.001, ^##^*p* < 0.01 and ^###^*p* < 0.001. OLB: olfactory bulb, PFC: prefrontal cortex, NAC: nucleus accumbens, STR: striatum, HIP: hippocampus, CEL: cerebellum.

Next, we assessed behavioral performances of *Glo1* KO mice to evaluate the effect of impairment of glyoxalase system on mouse behaviors. The *Glo1* KO mice showed shortened immobility time in the forced swimming test (FST) (**Figure 1H and 1I**) and less acoustic startle response in the prepulse inhibition (PPI) test without changes in PPI score (**Fig. 1J and 1K**).

However, the *Glo1* KO mice showed almost no changes in many behavioral tests, such as open-field test, social interaction tests, Y-maze test and novel object recognition test (**Suppl. Fig. S1**).

These findings suggest that gene deletion of *Glo1* alone is insufficient to cause schizophrenia-like behaviors.

### Combined effects of Glo1 KO and VB6 deficiency on methylglyoxal levels in the brain and mouse behaviors

Next, we sought to develop a novel *in vivo* model for MG detoxification deficits by an add-on treatment with an environmental factor, VB6 deficiency observed in more than 35% patients with schizophrenia^12–14^, to the *Glo1* KO mice. We fed the *Glo1* KO and WT mice a VB6-lacking diet containing low VB6 at 5 μg/100 g pellets from 8 to 12 weeks of age. We fed control mice a normal diet in which VB6 was present at 1.4 mg/100 g pellets. *Glo1* KO mice with the normal diet (KO/VB6(+)) showed lower plasma VB6 levels than WT mice with the normal diet (WT/VB6(+)), suggesting that *Glo1* deletion slightly decreases plasma VB6 levels. After feeding with the VB6-lacking diet for 4 weeks, we confirmed that plasma VB6 levels in WT (WT/VB6(-)) and KO (KO/VB6(-)) mice significantly decreased (**Fig. 2A**), although there is no change between them. Furthermore, feeding with the VB6-lacking diet decreased body weight in both WT and KO mice (**Fig. 2B**).

**Figure 2:**
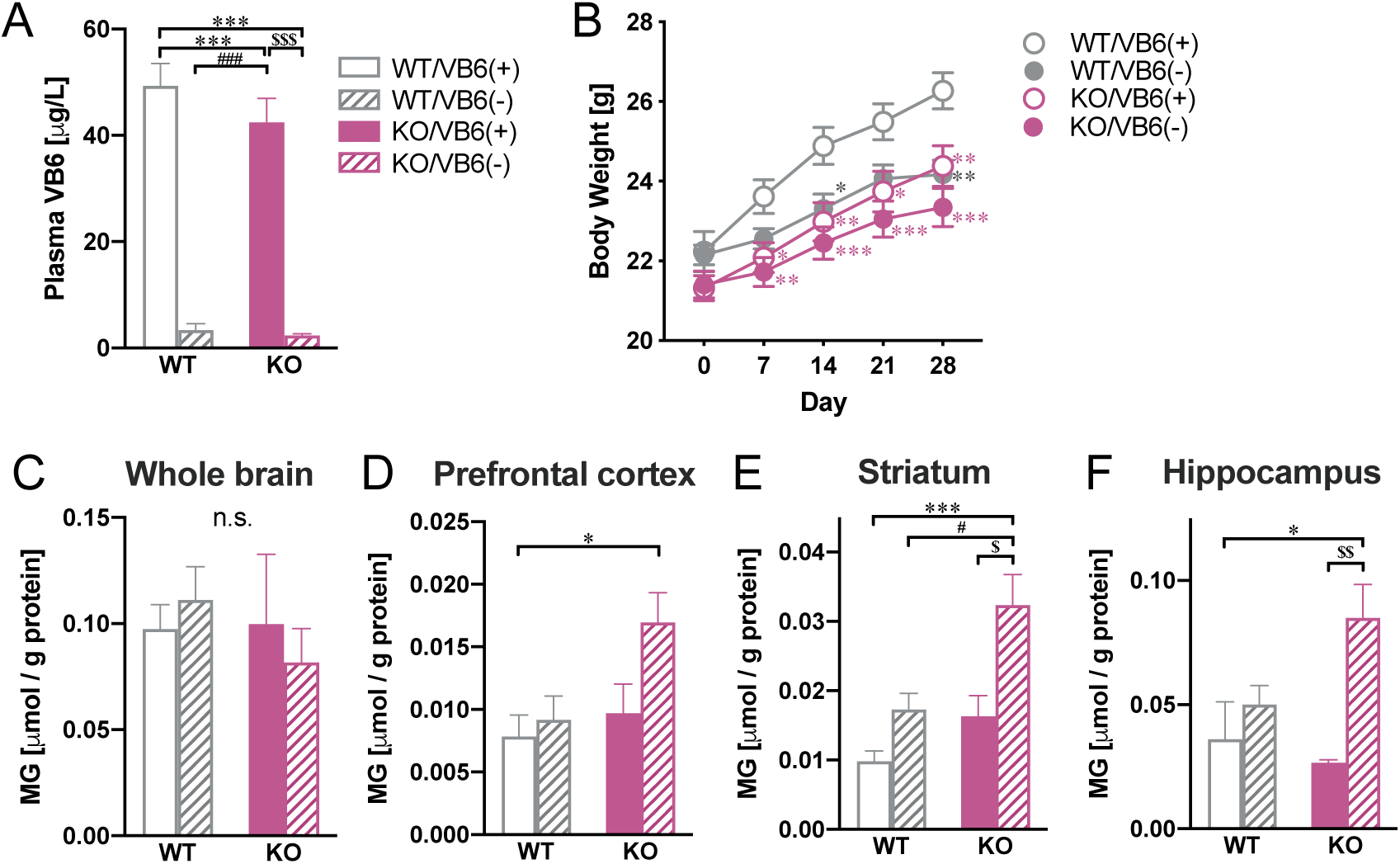
Development of Glo1 KO x VB6 deficiency mice. *Glo1* WT and KO mice were fed with VB6-lacking (VB6(-)) or control diet (VB6(+)) from 8 to 12 weeks. (A) VB6 levels in the plasma were determined (n = 27). (B) Changes in body weight during feeding with VB6(+) or VB6(-) diet are indicated (n = 14-18). Methylglyoxal levels were measured in (C) whole brain, (D) PFC, (E) STR and (F) HIP (n = 5). Two-way ANOVA: (A) *F*Interaction(1,104) = 0.87, *p >* 0.05; *F*Genotype(1,104) = 1.57, *p* > 0.05; *F*VB6(1,104) = 186, *p* < 0.0001, (B) *F*Interaction(12, 232) = 6.28; *p* < 0.0001, *F*Day(4, 232) = 212; *p*<0.0001, *F*Group(3, 58) = 5.53; *p* < 0.01, (C) *F*Interaction(1, 16) = 0.60; *p* > 0.05, *F*Genotype(1, 16) = 0.43; *p* < 0.05, *F*VB6(1, 16) = 0.01; *p* > 0.05, (D) *F*Interaction(1, 16) = 2.01; *p* > 0.05, *F*Genotype(1, 16) = 5.26; *p* < 0.05, *F*VB6(1, 16) = 4.19; *p* > 0.05, (E) *F*Interaction(1, 16) = 2.03; *p* > 0.05, *F*Genotype(1, 16) = 12.8; *p* < 0.01, *F*VB6(1, 16) = 15.2; *p* < 0.01, (F) *F*Interaction(1, 16) = 4.22; *p* > 0.05, *F*Genotype(1, 16) = 1.38; *p* > 0.05, *F*VB6(1, 16) = 11.2; *p* < 0.01. \**p* < 0.05, \*\**p* < 0.01, \*\*\**p* < 0.001, ^#^*p* < 0.05, ^###^*p* < 0.001, ^$^*p* < 0.05, ^$$^*p* < 0.01 and ^$$$^*p* < 0.001.

To evaluate the effect of MG detoxification deficits on the MG levels, we quantified MG in various brain regions of *Glo1* KO mice with VB6 deficiency. The MG levels in the prefrontal cortex (PFC: **Fig. 2D**), the striatum (STR: **Fig. 2E**) and the hippocampus (HIP: **Fig. 2F**) of the KO/VB6(-) group significantly increased compared to the WT/VB6(+) group, although there is no change in the whole brain (**Fig. 2C**) and other brain areas (**Suppl. Fig. S2**). There were no changes in MG levels in the WT/VB6(-) and KO/VB6(+) groups relative to the WT/VB6(+) group in all brain regions, suggesting that the accumulation of MG isn’t caused by *Glo1* gene deletion or VB6 deficiency alone, but by the combination of them.

Moreover, to investigate whether MG detoxification deficits affect the function of monoaminergic neuronal systems resulting in the behavioral impairments, we determined monoamines and their metabolites in various regions of the brain (**Suppl. Fig. S3**) and found that there were no changes in dopamine, noradrenaline (NA), or serotonin across all brain regions of KO/VB6(-) mice. We also demonstrated that 3-methoxy-4-hydroxyphenylglycol (MHPG) levels markedly increased in the VB6-deficient groups, WT/VB6(-) and KO/VB6(-), leading to enhanced NA turnover. This is in line with previous reports that VB6 deficiency induces the enhancement of NA metabolism (**Suppl. Fig. S3F and S3G**), although the differences were not significant.

Next, we assessed the behavioral performances of *Glo1* KO mice with VB6 deficiency. The VB6 deficient groups, WT/VB6(-) and KO/VB6(-), showed less locomotor activity in the first 10 minutes (**Fig. 3A**). In the novel object recognition test, the VB6 deficient groups also showed decreased exploratory preference during the retention session compared with the VB6 normal groups, although there was no change in exploratory time (**Fig. 3B and 3C**). In a social interaction test using a 3-chamber apparatus, the VB6 deficient groups spent less time in the chamber with an unfamiliar mouse and had decreased interaction time during the test session compared to the WT/VB6(+) group (**Fig. 3D and 3E**). These data demonstrate that the VB6-deficient mice display behavioral deficits of sociability and cognitive memory. Moreover, similar to the results in **Fig. 1H** to **1K**, the *Glo1* KO, KO/VB6(+), and KO/VB6(-) groups showed shortened immobility time in the FST (**Fig. 3F and 3G**) and less acoustic startle response in the PPI test (**Fig. 3H**). However, these behavioral abnormalities observed in the *Glo1* KO groups did not worsen with the combination of VB6 deficiency. Finally, the PPI deficit was only observed in the KO/VB6(-) group (**Fig. 3I**), suggesting that sensorimotor deficits occur only when the MG detoxification systems, glyoxalase system, and VB6-scavenge system are impaired.

**Figure 3:**
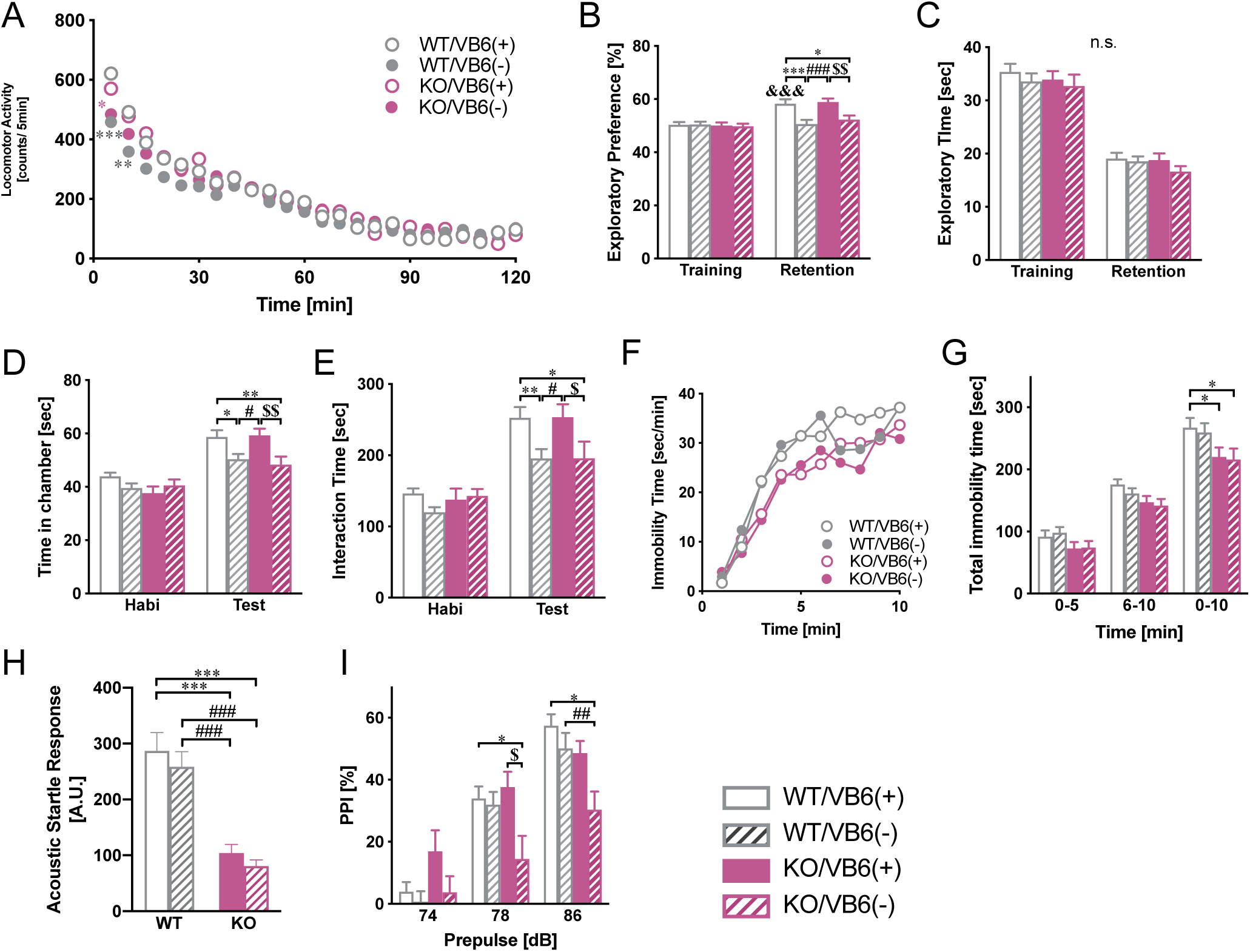
Behavioral deficits in Glo1 KO x VB6 deficient mice. (A) Time course of locomotor activity in the *Glo1* KO x VB6 deficiency mice were measured. (B) Exploratory preference and (C) exploratory time in the novel object recognition test, (D) time spent in the chamber and (E) interaction time in the social interaction test, (F) the time course and (G) total scores of the immobility time in the forced swimming test, and (H) the acoustic startle response without prepulse and (I) prepulse inhibition (PPI) score in the PPI test were measured. Two-way ANOVA: (A) *F*Interaction(69, 1863) = 1.56; *p* < 0.01, *F*Time(23, 1863) = 142; *p* < 0.0001, *F*Group(3, 81) = 0.39; *p* > 0.05, (B) *F*Interaction(3, 80) = 5.53; *p* < 0.01, *F*Session(1, 80) = 27.4; *p* < 0.0001, *F*Group(3, 80) = 5.00; *p* < 0.01, (C) *F*Interaction(3, 80) = 0.12; *p* > 0.05, *F*Session(1, 80) = 271; *p* < 0.0001, *F*Group(3, 80) = 1.00; *p* > 0.05, (D) *F*Interaction(3, 80) = 2.97; *p* < 0.05, *F*Session(1, 80) = 69.5; *p* < 0.0001, *F*Group(3, 80) = 5.19; *p* < 0.01, (E) *F*Interaction(3, 80) = 2.07; *p* > 0.05, *F*Session(1, 80) = 82.7; *p* < 0.0001, *F*Group(3, 80) = 4.40; *p* < 0.01, (F) *F*Interaction(27, 702) = 0.99; *p* > 0.05, *F*Time(9, 702) = 68.5; *p* < 0.0001, *F*Group(3, 78) = 2.47; *p* = 0.068, (G) *F*Interaction(6, 156) = 1.63; *p* > 0.05, *F*Genotype(2, 156) = 497; *p* < 0.0001, *F*VB6(3, 78) = 2.73; *p* < 0.05, (H) *F*Interaction(1, 79) = 0.01; *p* > 0.05, *F*Genotype(1, 79) = 47.3; *p* < 0.0001, *F*VB6(1, 79) = 0.98; *p* < 0.05, (I) *F*Interaction(6, 158) = 3.99; *p* < 0.01, *F*Prepulse(2, 158) = 136; *p*<0.0001, *F*Group(3, 79) = 3.94; *p* < 0.05. \**p* < 0.05, \*\**p* < 0.01, \*\*\**p* < 0.001, ^#^*p* < 0.05, ^##^*p* < 0.01, ^###^*p* < 0.001, ^$^*p* < 0.05 and ^$$^*p* < 0.01.

### Differential gene expression analysis in the brain of Glo1 KO x VB6-deficient mice

To identify molecular pathways impaired by *Glo1* KO and VB6 deficiency in the brain, we carried out RNA-seq in the PFC, STR, and HIP of the MG detoxification-impaired mouse model, where MG significantly accumulated (**Suppl. Table S1**). We identified only one differentially expressed gene (DEG), *Igf2*, in the PFC and STR, not HIP, by comparison between WT/VB6(+) vs WT/VB6(-), indicating the effect of the VB6 deficiency alone was minimal (**Fig. 4A**; **Suppl. Table S2**). We identified more DEGs by comparing WT/VB6(+) vs KO/VB6(+), indicating a greater effect of the *Glo1* deletion alone (**Fig. 4B**). Common DEGs in the three brain regions were *Glo1* and three zinc finger proteins (*Zfp40*, *Zfp758,* and *Zfp983*). Moreover, we identified many more DEGs in all brain regions by comparing WT/VB6(+) vs KO/VB6(-), reflecting the enhanced combined effect of the VB6 deficiency and the *Glo1* deletion (**Fig. 4C**). There were 10 common DEGs in the PFC, STR, and HIP (*Glo1, Zfp40, Atp6v0c, Zfp758, Pla2g7, Zfp983, Decr2, Rps2, Gnptg,* and *Zfp760*). It should be noted that the DEGs in the PFC were markedly increased to 286 genes compared with the other regions, suggesting that the combination of VB6 deficiency and the *Glo1* deletion has a greater impact on gene expression in the PFC.

**Figure 4:**
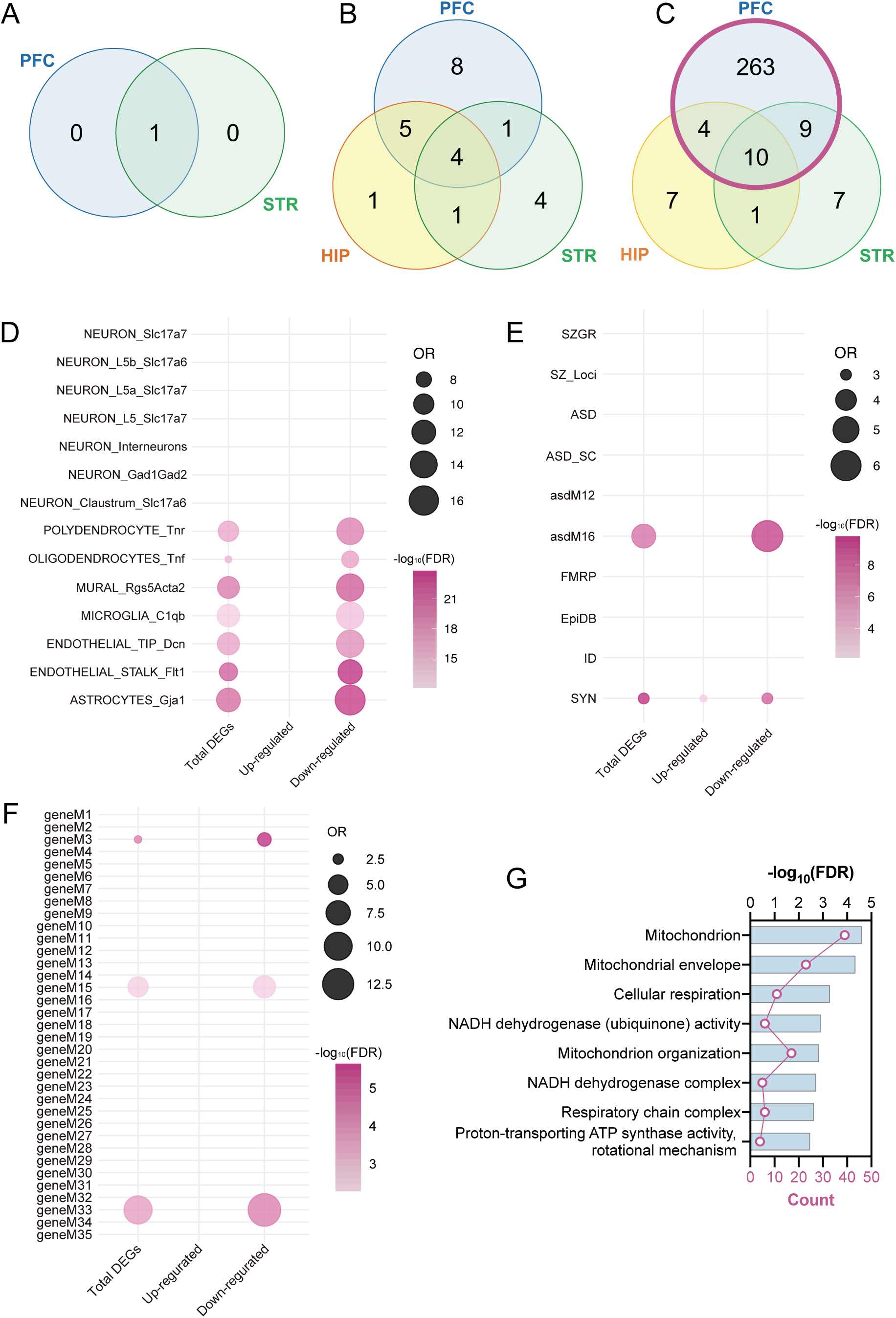
Changes of gene expression in the brain of Glo1 KO x VB6 deficiency mice. Venn diagrams of differentially expression genes (DEGs) overlapped among the PFC, STR, and HIP. (A) WT/VB6(-), (B) KO/VB6(+) and (C) KO/VB6(-) compared with WT/VB6(+) are shown. Gene set enrichment between DEGs in the PFC of KO/VB6(-) mice and (D) cell type markers from single-nuclei RNA-seq, (E) disease risk genes, or (F) modules associated with psychiatric disorders. Enrichment analyses were performed using a Fisher’s exact test. Odds ratios (OR) and -log10(FDR) are shown. (G) GO analyses of 216 down-regulated DEGs in the PFC between WT/VB6(+) and KO/VB6(-).

Cell type gene enrichment analysis indicated that the 286 DEGs in the PFC, 70 up-regulated and 216 down-regulated genes, were enriched for genes expressing specifically in endothelial cells and glial cells rather than neurons (**Fig. 4D; Suppl. Table S3**). The 286 DEGs were also enriched with genes in an asdM16 module, which is related to autism spectrum disorder (ASD), and enriched for genes involved in immune and inflammatory responses ^19^, but was not enriched for schizophrenia related genes (**Fig. 4E**). The enrichment results for the 286 DEGs are driven by the down-regulated genes, because the results of the down-regulated genes were the same as those of the 286 DEGs. To further examine the relationship between the PFC DEGs and dysregulation in neuropsychiatric disorders, used modules associated with schizophrenia, autism, or bipolar disorder^20^. We found that down-regulated genes are overrepresented for genes in a mitochondria-related module severely downregulated in schizophrenia (M33), an astrocyte module upregulated in autism and schizophrenia (M3), and a microglia module downregulated in schizophrenia (M15). These results further underscore the role of MG detoxification in schizophrenia etiologies and highlight the link between glial cells and neuropsychiatric disease etiologies (**Fig. 4F**).

Gene ontology (GO) analysis of the down-regulated DEGs identified enrichment for mitochondria-related function pathways, such as “Mitochondrial envelop,” “Mitochondrion,” and “Respiratory chain complex” (**Fig. 4G**), whereas that of the up-regulated genes enriched with “Whole membrane” only. Taken together, these findings suggest that the combination of VB6 deficiency and the *Glo1* deletion may impair gene expressions in endothelial and glial cells and induce mitochondrial dysfunction in the PFC.

### Weighted gene co-expression network analysis (WGCNA) and mitochondrial function in the PFC

To further identify gene co-expression modules in the PFC impaired by the combination of VB6 deficiency and *Glo1* deletion, we performed weighted gene co-expression network analysis (WGCNA)^21^ (**Fig. 5A; Suppl. Table S4**). We found two modules, “lightcyan” and “greenyellow”, that were associated with the combination of VB6 deficiency and *Glo1* deletion. The lightcyan module showed enrichment with DEGs in the PFC as well as mitochondria-related genes (**Fig. 5B-D**). GO analysis using the lightcyan module significantly identified enrichment for mitochondrial function (**Fig. 5E**). Considering that GO analysis using 286 DEGs in the PFC also identified enrichment for mitochondria-related functional pathways, these results suggest a hypothesis that altered gene expression may affect mitochondrial function in the PFC in the MG detoxification-impaired mouse. To test the hypothesis, we evaluated respiratory function of mitochondria isolated from the PFC in the *Glo1* KO mice with VB6 deficiency using a flux analyzer. We found that the oxygen consumption rates in the KO/VB6(-) mice were significantly decreased compared with the WT/VB6(+) (**Fig. 5F**). Together, these results indicate that the combination of VB6 deficiency and the *Glo1* deletion impaired mitochondrial respiratory function via aberrant gene expression in the PFC.

**Figure 5:**
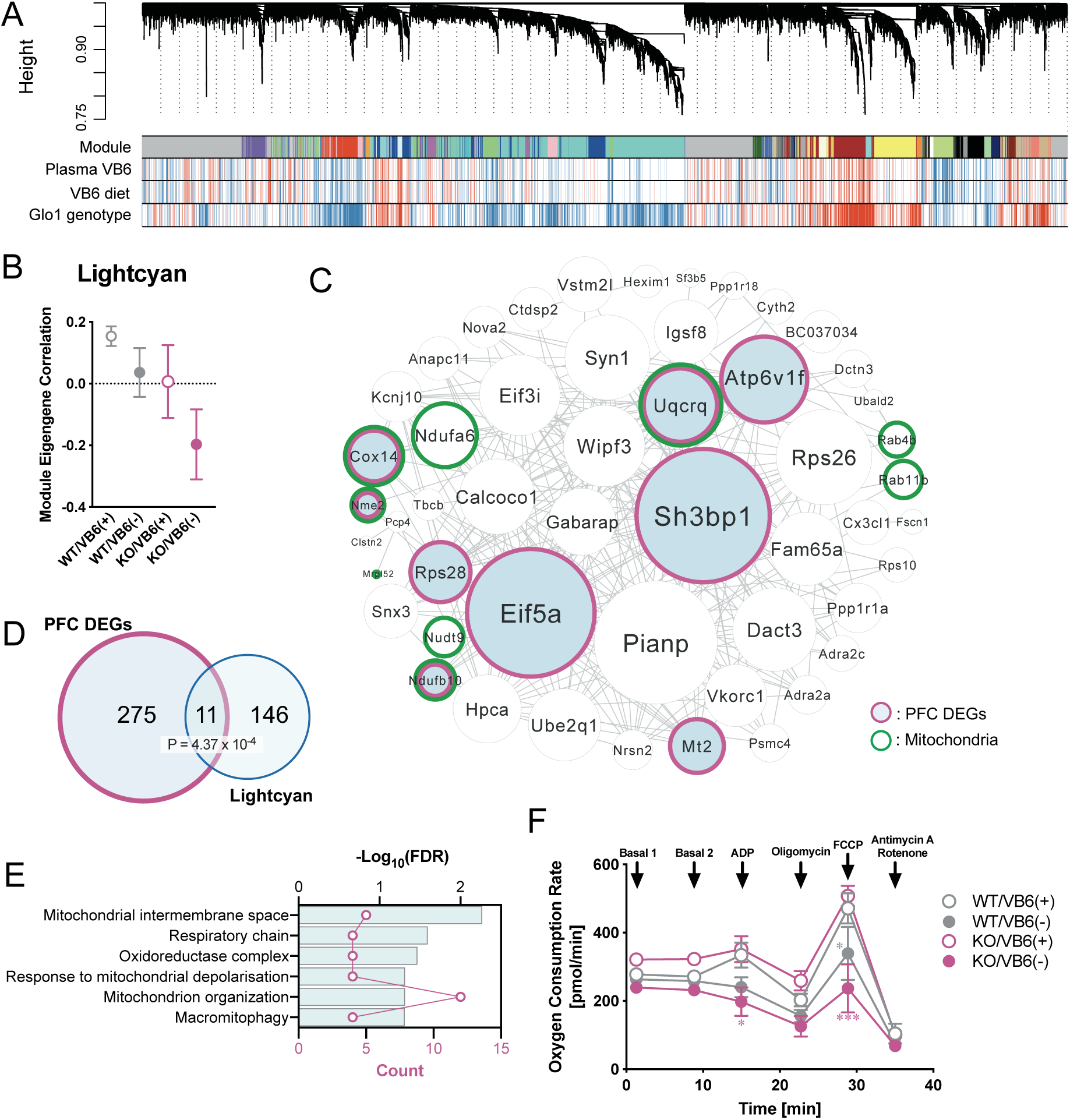
Weighted gene co-expression network analysis (WGCNA) and mitochondrial function in the PFC. (A) Representative network dendrograms. (B) Module eigengene (ME) of the lightcyan module significantly associated with the KO/VB6(-) group. SEs are calculated based on the eigengene across samples. Dots represent the mean eigengene for that module. (C) Visualization of the top 250 connections ranked by weighted topological overlap values for the lightcyan module. Node size corresponds to the number of edges (degree). DEGs in the PFC are highlighted in blue and surrounded by a purple frame. Mitochondria-related genes are surrounded by a green frame. (D) Venn diagrams of genes overlapped between the lightcyan module and DEGs in the PFC of KO/VB6(-) are shown [*p* = 4.37 x 10^-4^; hypergeometric test]. (E) GO analysis of genes in the lightcyan is shown. (F) Oxygen consumption rates of mitochondrial isolated from the PFC are measured by flux analyzer. Two-way ANOVA: (A) *F*Interaction(15, 80) = 2.11; *p* < 0.05, *F*Time(5, 80) = 54.0; *p* < 0.0001, *F*Group(3, 16) = 9.93; *p* < 0.001. \**p* < 0.05, and \*\*\**p* < 0.001.

## Discussion

In a previous report, we identified novel *GLO1* mutations thatreduce enzymatic activity in patients with schizophrenia^12^. To identify the mechanism of *GLO1* gene deletion on brain function, we first generated *Glo1* KO mice and evaluated behavioral phenotypes (**Fig. 1**). However, the *Glo1* KO mice showed few behavioral changes related to schizophrenia (**Fig. 1 and Suppl. Fig. S1**). Moreover, we also confirmed that the *Glo1* KO alone did not increase MG levels in the brain (Fig. 2C-2F). These findings imply that other compensatory systems remove MG in *Glo1* KO mice. For example, other enzymes such as aldehyde dehydrogenase and AKR metabolize MG to pyruvate and hydroxyacetone, respectively^22, 23^. Consistent with our findings, *in vitro* and *in vivo* studies show that in the absence of *Glo1*, AKR may compensate concerning MG detoxification, resulting in no MG accumulation^24, 25^. We also found no change in gene expression of AKR in the brain of *Glo1* KO mice (**Fig. 1C**). This suggests that the catabolic enzyme might compensate for the elimination of MG, which is usually removed by the glyoxalase system.

The *Glo1* KO mice exhibited shortened immobility time in the FST, a behavior associated with antidepressant-like behavior (**Fig. 1H and 1I**). This is consistent with a recent study that GLO1 inhibitors are useful as novel antidepressants^26^. Two structurally distinct GLO1 inhibitors decreased the immobility time in the FST and ameliorated the stress-induced depression-like behavior. Furthermore, the treatment with the GLO1 inhibitors increased the expression of molecular markers of the antidepressant response, including brain-derived neurotrophic factor (BDNF) and cyclic-AMP response-binding protein (CREB) phosphorylation in the HIP and the medial PFC. Considering our result that *Glo1* gene deletion alone is insufficient to increase MG levels in the brain, the therapeutic effect of the GLO1 inhibitors and the antidepressant-like behavior in the *Glo1* KO mice is probably independent of MG accumulation.

Next, to induce further impairments in MG detoxication systems, we depleted VB6 in the *Glo1* KO mice by feeding them VB6(-) diets. We found that the KO/VB6(-) mice displayed an accumulation of MG in the PFC, HIP, and STR (Fig. 2C-2F) as well as schizophrenia-like behavioral deficits, such as social deficits, cognitive impairment, and a sensorimotor deficit in the PPI test (**Fig. 3**). Moreover, RNA-seq expression analysis revealed a marked increase in DEGs in the PFC by the combination of *Glo1* gene deletion and VB6 deficiency (Fig. 2C-2F).

Gene enrichment analysis showed that the 286 DEGs in the PFC were enriched with genes expressing specifically in endothelial cells and glial cells, but not neurons, and overrepresented for genes in a mitochondria-related module downregulated in schizophrenia (**Fig. 4D and 4F**). MG is also produced in the periphery and delivered to the brain through the bloodstream; understandably, vascular endothelial cells are affected exogenously. Many studies have shown that MG can impair vascular endothelial cells in patients with diabetes and renal impairment^27^. In addition, astrocytes form the blood-brain barrier around blood vessels, so they are responsible for the supply of sugar from the bloodstream to neurons. This suggests that a disturbance in the MG detoxification system may enhance endogenous MG production in astrocytes. Therefore, it is also plausible that a disturbance in the MG detoxification system affects glial cells. Also, MG in the bloodstream appears to disrupt the blood-brain barrier, resulting in activation of the inflammatory response^28, 29^. Clinical studies have reported that the aberrant inflammation is frequently seen in schizophrenia and related psychoses^30, 31^. The fact that the 286 DEGs in the PFC are significantly correlated with AsdM16, which is associated with immune responses (**Fig. 4E**), and are also significantly enriched for immune-associated genes [GO:0002376, Immune system process, -log10 (p-value) = 1.38] suggests that inflammation may occur in the PFC of KO/VB6(-) mice. Furthermore, AGEs made from MG are also able to activate the immune response through receptor for AGEs (RAGE)^32^. Thus, AGEs produced from the accumulated MG in the PFC of KO/VB6(-) mice may lead to the activation of inflammation.

An association between PPI deficits observed in KO/VB6(-) mice and glial cell damage may fit with previous reports. For example, the expression of astrocytic markers is variable in the PFC of a mouse model for schizophrenia showing PPI impairment^33^. Cortical myelination by oligodendrocytes correlates with PPI deficits in a mouse model for schizophrenia^34^. Furthermore, chimeric mice of glia cells differentiated from induced pluripotent stem (iPS) cells derived from patients with schizophrenia displayed PPI deficits^35^. These findings suggest that glial cells may be impaired in schizophrenic patients with impaired MG detoxification systems.

We also found that VB6 deficiency alone induced social deficits and cognitive impairment (**Fig. 3A-3E**). We have already linked behavioral deficits induced by VB6 deficiency to enhanced noradrenergic signaling (Toriumi et al. *Submitted*). The VB6-deficient groups WT/VB6(-) and KO/VB6(-) showed a marked increase in MHPG, a metabolite of noradrenaline, in the brain (**Suppl. Fig. 3**), although the combination of VB6 deficiency and *Glo1* gene deletion did not increase MHPG synergistically. However, like *Glo1* gene deletion, VB6 deficiency alone is not enough to increase MG levels in the brain (**Fig. 2C-2F**). Thus, the noradrenergic system activation by VB6 deficiency is not due to MG accumulation in the brain. To identify molecular mechanisms underlying our observed finding that VB6 deficiency increases noradrenergic signaling, further experiments are required.

Finally, we performed WGCNA analysis and found that the gene co-expression module “lightcyan” associated with mitochondrial function in the PFC of KO/VB6(-) mice (Fig. 5A-E). Since enrichment and GO analysis using DEGs also revealed mitochondrial dysfunction (**Fig. 4F and 4G**), we isolated mitochondria from the PFC of KO/VB6(-) mice and found that oxidative phosphorylation was impaired (**Fig. 5F**). Mitochondrial dysfunction has been frequently reported in patients with schizophrenia^36^. Several studies using postmortem brains demonstrated a decrease in the activity of complex IV in the PFC of patients with schizophrenia^37^, and a global down-regulation of mitochondria-related genes by microarray analysis^38^. Also, mitochondrial dysfunction has also been found in patient-derived iPS cells^39, 40^; MG can disrupt mitochondrial respiration^41^. Incubation of isolated mitochondria with MG produced a concentration-dependent decrease in state III as well as an increase and then a decrease in state IV respiration. These findings suggest that the accumulated MG may affect mitochondrial function in the KO/VB6(-) mice, causing schizophrenia-like sensorimotor deficits. Additionally, the disturbance of the respiratory chain leads to the production of ROS and further production of MG. The aberrant gene expression related to the mitochondria functional pathway in the PFC may contribute to further MG accumulation.

Another module from the WGCNA, the “greenyellow” module, was also associated with the combination of VB6 deficiency and *Glo1* deletion (**Suppl. Fig. S4A and S4B**). GO analysis of the greenyellow module identified significant enrichment for RNA splicing and microtubule-related pathways (**Suppl. Fig. S4C**). Thus, we checked differential exon usage using the DEXseq method in our RNA-seq data^42^ and identified genes with differential exon usage in the PFC of *Glo1* KO mice with VB6 deficiency (**Suppl. Table S5).** We also found that the genes with differential exon usage were enriched for synaptic function by GO analysis (**Suppl. Fig. S4C**). Next, to evaluate synaptic function, we carried out an electrophysiological analysis of the PFC of *Glo1* KO mice with VB6 deficiency. However, the results showed no significant differences in both miniature excitatory postsynaptic current (mEPSC) and miniature inhibitory postsynaptic current (mIPSC) among all groups, although we observed increased mEPSC amplitude in the KO/VB6(+) group and depolarized membrane potential in the KO/VB6(-) (**Suppl. Fig. S4E-S4K**). These findings suggest that the effect of alternative splicing on synaptic function in the PFC of *Glo1* KO mice with VB6 deficiency might be minimal.

Although it remains unclear how MG accumulation connects to altered gene expression in the PFC of KO/VB6(-) mice, recent studies provide a possible answer to this question: MG-derived modification of histones and DNA. Recent reports claim that MG causes glycative modification of histones involved in the regulation of gene expression^43, 44^. DJ-1, an enzyme that removes MG-modification of histones, has also been identified, and this MG-modification is physiologically regulated in cancer cells. Also, this MG-mediated glycative modification also occurs in DNA^45^, and DJ-1 can also remove MG-derived modifications on DNA^46^, suggesting that MG-derived modification of DNA may also be involved in the regulation of regulatory gene expression. Further study is necessary to understand molecular mechanisms underlying the altered gene expression in KO/VB6(-) mice.

Taken together, we generated *Glo1* KO mice to evaluate the effect of MG detoxification deficits on brain function and found that *Glo1* KO mice fed VB6(-) diets exhibited an accumulation of MG in the PFC, STR, and HIP as well as sensorimotor deficits. Furthermore, we found altered gene expression involved in mitochondria function in the PFC of the KO/VB6(-) mice and we demonstrated respiratory deficits in mitochondria of their PFC.

## Methods

### Generation of Glo1 knockout mice

A 129/Sv-derived ES cell line (S17-2B1) containing a gene trap cassette in the *Glo1* gene locus was purchased from Mutant Mouse Regional Resource Centers (http://www.mmrrc.org/). The ES cells were microinjected into embryonic day 3.5 (E3.5) blastocysts (genotype C57BL/6J) and then transferred into pseudopregnant ICR females for the generation of chimeras. Chimerism of newborn mice was assessed by coat color inspection. A male chimera with high ES cell contribution (100% agouti coat color) was crossed with C57BL/6J mice. The germline transmission was judged by the presence of agouti pups (F1 generation) and disrupted alleles identified by PCR genotyping as indicated below.

One pair of heterozygous (HE) F1 mice were mated to confirm whether the generated F2 *Glo1* KO mouse was nonlethal and normal judging by appearances or not. One randomly selected HE F1 male mouse was backcrossed onto C57BL/6J female mice. The procedure was repeated for ten generations. The HE F11 mice were intercrossed and the behavioral tests were performed with the F12 generation. KO, HE, and wild-type mice (WT) were identified by PCR genotyping before use.

### PCR Genotyping

Three primers were designed from the sequences of the *Glo1* locus and gene trap cassette. The presence of the wild-type allele (200 bp) or gene-trap allele (284 bp) was detected with primer pairs GLO1F (5’-GAG ACC GAG ACA CAC CCC TA-5’) and GLO1R (5’-ATG TGT TCG CTG TGC ACC TA-3’) or GLO1F and GEOR (5’-GGT CCA GGC TCT AGT TTT GAC TC-3’), respectively, using tail genomic DNA. PCR cycles were as follows: initial denaturation at 98 °C for 2 min, 35 cycles at 98 °C for 10 sec, 60 °C for 15 sec, and 68 °C for 1 min. PCR products were separated in 2% (w/v) agarose gel and stained with ethidium bromide.

### Glo1 mRNA expression analysis

Total RNAs were extracted from the brains of KO, HE or WT mice using the Fast Pure RNA kit (Takara). The purified RNA samples were used as templates for reverse transcription reactions. Qualitative PCR assays for wild-type transcripts of *Glo1* or fusion transcripts of *Glo1* exon 1 and βGEO were performed. Quantitative PCR for *Glo1* was performed with iQ SYBR Green Supermix and amplified PCR products were measured by the iCycler iQ real-time PCR analyzing system (Bio-Rad).

### GLO1 activity assay

GLO1 enzymatic activity was determined using the spectrophotometric method as described previously with minor modifications^47^. Whole brains were homogenized in a complete Lysis-M reagent (Sigma-Aldrich) containing a protease inhibitor cocktail (cOmplete: Sigma-Aldrich) and a phosphatase inhibitor (PhosSTOP: Sigma-Aldrich). After centrifugation at 15,000 rpm for 15 min at 4°C, the supernatants were collected and diluted to 1 mg/ml in the Lysis-M reagent. The substrate of *GLO1*, hemithioacetal, was prepared by preincubation of 2 mM methylglyoxal with 2 mM glutathione in a 50 mM sodium phosphate buffer (pH 6.6) at 37 °C for 10 minutes. The supernatants of the brain (4 μl) were mixed with the hemithioacetal solution (196 μl), and displayed an absorbance at 240 nm, owing to the formation of S-D-lactoylglutathione as measured for 15 min by a microplate spectrophotometer EPOCH2 (BioTek).

### Forced Swimming Test

Each mouse was placed in an opaque polyvinyl bucket (24 cm high, 23 cm in diameter), which contained water at 22–23 °C to a depth of 15 cm, and was forced to swim for 10 min. The duration of swimming was measured using an infrared detector (Muromachi Kikai Co. Ltd., Japan). The duration of swimming was measured every minute by an infrared detector.

Immobility time was calculated as follows: total time (sec) – swimming time (sec) = immobility time (sec).

### Prepulse inhibition (PPI) test

PPI of the acoustic startle response was measured using an SR-LAB System (San Diego Instruments). The stimulus consisted of a 20-ms prepulse, a 100-ms delay, and then a 40-ms startle pulse. The intensity of the prepulse was 4, 8, or 16-dB above the 70-dB background noise. The amount of PPI was calculated as a percentage of the 120-dB acoustic startle response: 100 – [(startle reactivity on prepulse + startle pulse)/startle reactivity on startle pulse] x 100.

### Locomotor activity test

Locomotor activity was measured using an infrared detector (Muromachi Kikai Co. Ltd., Japan). Mice were placed in a plastic round arena (27cm high, 27 cm in diameter) and locomotor activity was measured over a period of 2 hours.

### Novel object recognition test

The novel object recognition test was performed as described previously with minor modifications^48^. The test procedure consisted of three sessions: habituation, training, and retention. Each mouse was individually habituated to the box, with 10 min of exploration in the absence of objects each day for 3 consecutive days (Day 1-3) (habituation session). On day 4, each animal was allowed to explore the box for 10 min, in which two novel objects were placed symmetrically. The time spent exploring each object was recorded (training session). The objects were different in shape and color but similar in size. Mice were considered to be exploring an object when their heads were facing it or when they were sniffing it at a distance of less than 2 cm and/or touching it with their nose. After the training session, mice were immediately returned to their home cages. On day 5, the animals were placed back into the same box with one of the familiar objects used in the training session and one novel object. Mice were allowed to explore freely for 5 min and the time spent exploring each object was recorded (retention session). An exploratory preference, the ratio of time spent exploring either of the two objects (training session) or the novel object (retention session) over the total amount of time spent exploring both objects, was used to assess recognition memory.

### Social interaction test

Twelve-week-old male mice were tested for social behavior using the three-chamber social approach paradigm. The apparatus was a square field (W50 × D50 × H40 cm) that is divided into three chambers of equal size by grey walls, which was placed in a dark, sound-attenuated room. During a habituation session, two empty wire cages were placed in the lateral chambers. The mice were placed in the center chamber and allowed to freely explore all chambers for 10 min. Immediately after the habituation session, the mouse was placed in a home cage for 3 min. After the interval, a test session started. The mice were placed in the center chamber again and allowed to freely explore the left and right arenas, which contained a social target (unfamiliar C57BL/6J male mouse) in one chamber and empty in the other chamber. Experimental mice were given 10 min to explore both chambers. The amount of time spent in the lateral chambers and the quadrant around each wire cage as interaction time (that is, sniffing, approach) was recorded and analyzed by the EthoVision tracking system (Noldus, Netherland).

### Induction of VB6 deficiency

To induce VB6 deficiency, WT and KO mice were fed with a VB6-lacking diet containing 5 μg/100 g VB6 pellets from 8 to 12 weeks of age, while control mice in normal VB6 condition were fed with a normal diet, with 1.4 mg/100 g VB6 pellets. Food and tap water were available *ad libitum*.

### Measurement of VB6 in mouse plasma

VB6 concentration in plasma was determined with ID-Vit VB6 (Immunodiagnostic AG: Bensheim, Germany) according to the manufacturer’s protocol.

### Measurement of methylglyoxal in the brain

Methylglyoxal in the brain was quantified using high-performance liquid chromatography with fluorescence (HPLC-FL) as described previously^49, 50^. Brain tissues were homogenized using BioMasher (Nippi, Japan) with nine volumes of ice-cold buffer: 0.1M phosphate buffer [pH 7.8] and a protease inhibitor cocktail (Nacalai Tesque, Japan). The brain extracts were centrifuged at 12,000 × g for 10 min at 4 °C. Subsequently, trichloroacetic acid was added to a final concentration of 4% and centrifuged at 12,000 × g for 10 min at 4 °C. The supernatant was diluted with an equal volume of 1,2-diamino-4,5-methylenedioxybenzene (MDB) solution and incubated at 60 °C for 40 min in the dark. This MDB-derived sample was then analyzed using HPLC-FL. Ten microliters of the resulting mixture were injected into a C18 column pre-equilibrated with a mobile phase solution, which consisted of water, methanol, and acetonitrile in the ratio 10:7:3. A flow rate of 1.0 ml/min was used with a running time of 7 min. The retention times and peak areas were monitored at excitation and emission wavelength of 355 nm and 393 nm, respectively.

### RNA sequencing

RNA-sequencing (RNA-seq) was performed as described previously^51^. Briefly, total RNA from the PFC, STR, and HIP was extracted with the miRNeasy Mini Kit (Qiagen, 217004) according to the manufacturer’s instructions. RNA integrity number (RIN) of total RNA was quantified by an Agilent 2100 Bioanalyzer using an Agilent RNA 6000 Nano Kit (Agilent, 5067-1511). Total RNAs with an average RIN value of 8.52 ± 0.05 were used for RNA-seq library preparation. mRNA was purified from 2 μg of total RNA by NEXTflex poly(A) beads (Bioo Scientific, 512981), subjected to fragmentation and first and second strand syntheses, and cleaned up by EpiNext beads (EpiGentek, P1063). The second strand DNA was adenylated, ligated, and cleaned up twice by EpiNext beads. cDNA libraries were amplified by PCR and cleaned up twice by EpiNext beads. cDNA library quality was quantified by a 2100 Bioanalyzer using an Agilent high-sensitivity DNA kit (Agilent, 5067-4626). Libraries were sequenced separately as 75-bp single ends on an Illumina NextSeq 500.

### RNA-seq data processing

An in-house RNA-seq pipeline was established for RNA-seq data analysis and processing. Reads were aligned to the mouse mm10 reference genome using STAR 2.5.2b^52^ with the following parameter --outFilterMultimapNmax 10 --alignSJoverhangMin 10 --alignSJDBoverhangMin 1 -- outFilterMismatchNmax 3 --twopassMode Basic. Gencode annotation for mm10 (version vM11) was used as a reference to build STAR indexes and alignment annotation^53^. For each sample, a BAM file including mapped and unmapped with spanning splice junctions was produced. Secondary alignment and multi-mapped reads where further removed using in-house scripts. Only uniquely mapped reads were retained for further analysis.

Overall quality control metrics were performed using RseqQC using the UCSC mm10 gene model provided^54^. This includes the number of reads after multiple-step filtering, ribosomal RNA reads depletion, reads mapped to an exon, UTRs, and intronic regions.

Gene level expression was calculated using HTseq version 0.6.0 using intersection-strict mode by exonic regions^55^. Counts were calculated based on protein-coding genes annotation from mm10 Gencode gtf file (version vM11). CPM (counts per million reads) was calculated using edgeR^56^. To retain only expressed genes, we used a “by condition” log2(CPM + 1) cutoff. Briefly, a gene is considered expressed with log2(CPM + 1) >= 0.5 in all four replicates of a given condition (WT/VB6(+), WT/VB6(-), KO/VB6(+), KO/VB6(-)). In total, we detected 13428 protein-coding genes expressed in our data. We further used those genes for differential and co-expression analyses.

### Differential expression analysis

Differential expression was assessed using the DESeq2 package in R^57^. We applied the following method on all the pairwise comparisons between WT and KO, with/without VB6 in the diet.

The expression matrix contained the 13428 protein-coding genes that passed the log2(CPM + 1) >= 0.5 cut-off in all four replicates of an experimental condition and the covariate matrix including RIN values and Batches were included.

Using the SVA package in R^58^, we identified potential unwanted surrogate variables (2 surrogate variables).

Linear regression was performed using DESeq2 to remove covariate variables: Gene expression ∼ SVA1 + SVA2 + Batch We calculated differentially expressed genes (DEGs) using criteria of FDR <= 0.05 and log2(fold change) >= |0.3|. A permutation test was additionally applied using 1000 permuted experiments. None of the permuted analyses showed the same genes differentially expressed (permutation *p* < 0.001).

Gene ontology (GO) analysis of the significant DEGs was carried out using ToppGene (https://toppgene.cchmc.org), and the resultant GO terms were consolidated using REVIGO (reduce and visualize GO)^59^.

### Co-expression network analysis (WGCNA)

To identify the modules of co-expressed genes, we carried out weighted gene co-expression network analysis (WGCNA)^21^ on the RNA-seq data. As input, we used log2(CPM+1) values of the 13428 protein-coding expressed genes. Batch effects were further removed using the ComBat function in the SVA package in R^58^. We generated a signed network by using the blockwiseModules function in the WGCNA package. Beta was chosen as 14 so the network has a high scale-free R square (r^2^ = 0.79). For other parameters, we used corType = bicor, maxBlockSize = 14000, mergingThresh = 0.10, reassignThreshold = 1e-5, deepSplit = 4, detectCutHeight = 0.999, and minModuleSize = 50. Then the modules were determined by using the dynamic tree-cutting algorithm.

The top 250 edges based on ranked weight were plotted to represent the modules showing the top hub genes. We used Cytoscape 3.7.2 for data visualization and optimization^60^.

### Identification of cell type preferential expression

To identify cell type preferential expression of the DEGs and detected modules, we used a scRNA-seq data from mouse brain curated from DropViz database^61^. Cell-types markers were downloaded and used for the enrichment test. We used a Fisher’s exact test in R with the following parameters: alternative = “greater”, conf.level = 0.99. We reported Odds Ratio (OR) and Benjamini-Hochberg adjusted *P*-values (FDR). This method was used for both DEGs and the WGCNA module enrichment described below.

### Collected data sources

Only genes that are expressed in the mouse brain samples are included as the background information. The ASD-related genes were obtained from the Simons Foundation Autism Research Initiative^62^. ASD modules were downloaded from an independent publication^19^. Intellectual disability genes were collected from multiple independent sources^63^. Genes related to synapse were downloaded from the SynaptomeDB^64^. Schizophrenia associated genes were curated from the SZGR database^65^. For fragile X mental retardation (FMRP) gene targets, we used an independent publication^66^. Epilepsy associated genes were curated from the Epilepsy Gene Database^67^. scRNA-seq from mouse brain was curated from DropViz databse^61^. Modules associated with neuropsychiatric disorders were curated from an independent publication^20^. All genes from human collections were translated into mouse gene IDs using the biomaRt package in _R_^68^.

### Measurement of respiratory function in isolated PFC mitochondria

Mitochondria were isolated from the PFC with a mitochondria isolation kit for tissue and cultured cells (Biochain Institute, CA) according to the manufacturer’s protocol.

Respiratory function in the isolated mitochondria was quantified using extracellular Flux Analyzer Seahorse XFe96 (Agilent technologies) as described previously with minor modifications^69, 70^. Isolated mitochondria were diluted at 0.2 mg/ml with mitochondria assay solution [70mM sucrose, 210 mM mannitol, 2 mM HEPES, 5 mM MgCl2, 10m M KH2PO4, 0.2 % fatty acid-free BSA, 10 mM sodium pyruvate, 2 mM D-malic acid (pH 7.4)]. The mitochondria (100 μl) were transferred into a well of an assay plate and attached on the bottom of the well by centrifugation at 2,000 × g for 20 min at 4 °C. After incubation at 37 °C for 10 min, the measurement of oxygen consumption rate was calculated according to the manufacturer’s protocol. Injections were as follows : port A, 20 μl of 30 mM ADP (5 mM final); port B, 20 μl of 35 μM oligomycin (5 μM final); port C, 20 μl of 40 μM FCCP (5 μM final); and port D, 20 μl of 90 μM antimycin A (10 μM final) and 18 μM antimycin A (18 μM final).

Typical mix and measurement cycle times for the assay were the same as previous reports ^69, 70^.

### Statistical Analysis

In the behavioral and biochemical studies, statistical differences between the two groups were determined with a Student’s *t*-test. Statistical differences between more than 3 groups were determined using a one-way or two-way ANOVA with repeated measures followed by Tukey’s multiple comparisons test. A value of *p* < 0.05 was considered statistically significant.

### Study approval

For all animal studies, the experimental procedures were approved by the Animal Experiment Committee of the Tokyo Metropolitan Institute of Medical Science (approval no. 15001, 16001, 17003, 18006, 19003, 20005).

## Data Availability

RNA-seq data are available in the NCBI Gene Expression Omnibus (GEO): accession number is GSE149585 (token: crwvwyysvnipxwz). Also, Custom R codes and data to support the analysis, visualizations, functional and gene set enrichments are available at: https://github.com/konopkalab/Glo1_Vitamin

## Acknowledgments

We thank the donors and their families for the samples used in these studies. We are grateful for the expert technical assistance of Izumi Nohara, Yukiko Shimada, Emiko Hama, Nanako Obata, Mai Hatakenaka, Chikako Ishida, and Ikuyo Kito. We would also like to express my gratitude to Connor Douglas for his proofreading supports. This work was supported by the JSPS Program for Advancing Strategic International Networks to Accelerate the Circulation of Talented Researchers (S2603); JSPS KAKENHI Grant Numbers 16H05380, 17H05930, 18K06977, 19H04887, 20H03608; AMED under Grant Number JP20dm0107088; the Kanae Foundation for the Promotion of Medical Science; The Uehara Memorial Foundation; The Sumitomo Foundation.

## Author contributions

K.T. designed and conducted the study and wrote the initial draft of the manuscript. S.B. and N.U. conducted RNA-seq analysis. S.K. and Y.O. quantified MG in the brain. T.D. and T.M. generated *Glo1* KO mice. K.S., M.M., Y.H., A.Y. and Mai Asakura. provided technical support. Y.S., T.W., and M.K. performed electrophysiological analysis. Makoto Arai, G.K., and M.I. supervised the study and provided intellectual guidance. All authors discussed the results and commented on the manuscript.

## Competing interests

The authors have declared that no conflict of interest exists.

## Notes

### Competing Interest Statement

The authors have declared no competing interest.

