## Supplemental Figures for "Impairment of methylglyoxal detoxification systems causes mitochondrial dysfunction and schizophrenia-like behavioral deficits"

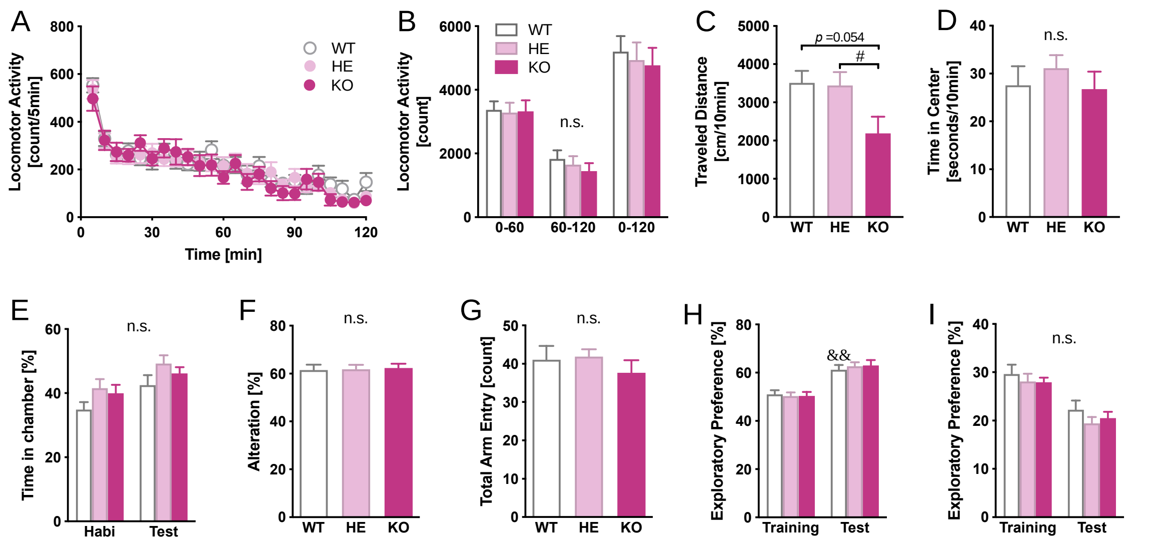


***Supplementary Figure 1: Behavioral phenotypes of Glo1 knockout mice***

In the locomotor activity test, (A) time course and (B) total locomotor activity were determined. Two-way ANOVA: (A) $F_{\text{Interaction}(46,1035)} = 0.88$, $p > 0.05$; $F_{\text{Time}(23,1035)} = 47.6$, $p < 0.0001$; $F_{\text{Genotype}(2, 45)} = 0.14$, $p > 0.05$, (B) $F_{\text{Interaction}(4,90)} = 0.30$, $p > 0.05$; $F_{\text{Time}(2,90)} = 234$, $p < 0.0001$; $F_{\text{Genotype}(2, 45)} = 0.14$, $p > 0.05$. In the open field test, (C) traveled distance and (D) time spent in the center area were measured. One-way ANOVA: (C) $F_{(2, 44)} = 3.59$, $p < 0.05$, $^{\#}p < 0.05$, (D) $F_{(2, 44)} = 0.53$, $p > 0.05$. (E) In the social interaction test, time spent in the chamber was measured. Two-way ANOVA: $F_{\text{Interaction}(2, 48)} = 0.11$; $p > 0.05$, $F_{\text{Session}(1, 48)} = 20.1$; $p < 0.0001$, $F_{\text{Genotype}(2, 48)} = 2.14$; $p > 0.05$. In the Y-maze test, (F) alteration and (G) total arm entry were determined. One-way ANOVA: (F) $F_{(2, 45)} = 0.04$, $p > 0.05$, (G) $F_{(2, 45)} = 0.63$, $p > 0.05$. In the novel object recognition test, (H) exploratory preference and (I) exploratory time were measured. Two-way ANOVA: (H) $F_{\text{Interaction}(2, 42)} = 0.28$; $p > 0.05$, $F_{\text{Session}(1, 42)} = 75.2$; $p < 0.0001$, $F_{\text{Genotype}(2, 42)} = 0.04$; $p > 0.05$, $^{\&\&}p < 0.01$ (Training vs. Test in WT mice), (I) $F_{\text{Interaction}(2, 42)} = 0.15$; $p > 0.05$, $F_{\text{Session}(1, 42)} = 43.7$; $p < 0.0001$, $F_{\text{Genotype}(2, 42)} = 0.90$. The data are represented as the mean ± SEM values.

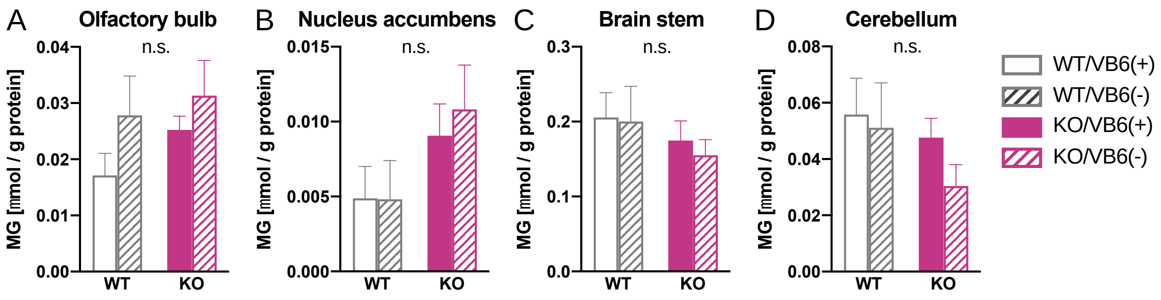


***Supplementary Figure 2: Methylglyoxal in Glo1 KO x VB6 deficiency mice***

Methylglyoxal level were measured in (A) olfactory bulb, (B) nucleus accumbens, (C) brain stem and (D) cerebellum (n = 5). Two-way ANOVA: (A) $F_{\text{Interaction}(1,16)} = 0.19$, $p > 0.05$; $F_{\text{Genotype}(1,16)} = 1.24$, $p > 0.05$; $F_{\text{VB6}(1,16)} = 2.58$, $p > 0.05$, (B) $F_{\text{Interaction}(1,16)} = 0.13$, $p > 0.05$; $F_{\text{Genotype}(1,16)} = 4.26$, $p > 0.05$; $F_{\text{VB6}(1,16)} = 0.12$, $p > 0.05$, (C) $F_{\text{Interaction}(1,16)} = 0.04$, $p > 0.05$; $F_{\text{Genotype}(1,16)} = 1.31$, $p > 0.05$; $F_{\text{VB6}(1,16)} = 0.14$, $p > 0.05$, (D) $F_{\text{Interaction}(1,16)} = 0.30$, $p > 0.05$; $F_{\text{Genotype}(1,16)} = 1.60$, $p > 0.05$; $F_{\text{VB6}(1,16)} = 0.92$, $p > 0.05$. The data are represented as the mean ± SEM values.

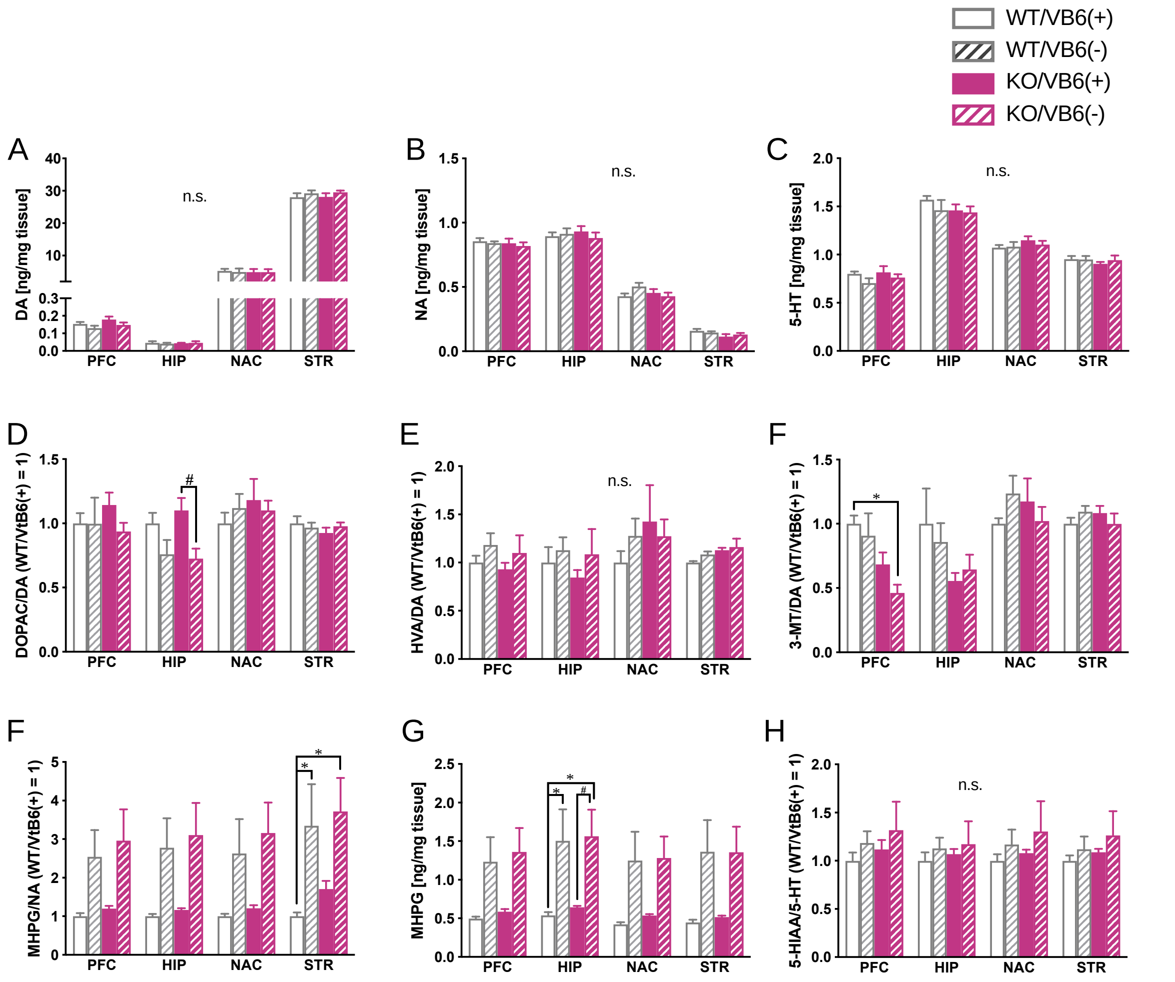


***Supplementary Figure 3: Monoamine contents in the brain of Glo1 KO x VB6 deficiency mice***

(A) dopamine (DA), (B) noradrenaline (NA), and (C) serotonin (5-HT: 5-hydroxytryptamine) contents, (D - F) DA, (G) NA and (I) 5-HT turnover and (H) MHPG contents in various regions of the mouse brain were determined. Two-way ANOVA: (A) $F_{\text{Interaction}(9,60)} = 0.39$, $p > 0.05$; $F_{\text{Region}(3, 60)} = 1780$, $p < 0.0001$; $F_{\text{Group}(3, 20)} = 0.20$, $p > 0.05$, (B) $F_{\text{Interaction}(9,60)} = 0.71$, $p > 0.05$; $F_{\text{Region}(3, 60)} = 750$, $p < 0.0001$; $F_{\text{Group }(3, 20)} = 0.79$, $p > 0.05$, (C) $F_{\text{Interaction}(9,60)} = 1.76$, $p > 0.05$; $F_{\text{Region}(3, 60)} = 339$, $p < 0.0001$; $F_{\text{Group }(3, 20)} = 0.29$, $p > 0.05$, (D) $F_{\text{Interaction}(9,60)} = 1.37$, $p > 0.05$; $F_{\text{Region}(3, 60)} = 4.05$, $p < 0.05$; $F_{\text{Group }(3, 20)} = 1.06$, $p > 0.05$, (E) $F_{\text{Interaction}(9,60)} = 0.62$, $p > 0.05$; $F_{\text{Region}(3, 60)} = 1.86$, $p > 0.05$; $F_{\text{Group}(3, 20)} = 0.66$, $p$

> 0.05, (F) $F_{Interaction(9,60)} = 1.92$, $p > 0.05$; $F_{Region(3, 60)} = 12.0$, $p < 0.0001$; $F_{Group\ (3, 20)} = 1.90$, $p > 0.05$, (G) $F_{Interaction(9,60)} = 1.77$, $p > 0.05$; $F_{Region(3, 60)} = 13.5$, $p < 0.0001$; $F_{Group\ (3, 20)} = 3.45$, $p < 0.05$, (H) $F_{Interaction(9,60)} = 3.11$, $p < 0.05$; $F_{Region(3, 60)} = 28.1$, $p < 0.0001$; $F_{Group\ (3, 20)} = 3.94$, $p < 0.05$, (I) $F_{Interaction(9,60)} = 0.49$, $p > 0.05$; $F_{Region(3, 60)} = 1.57$, $p > 0.05$; $F_{Group\ (3, 20)} = 0.51$, $p > 0.05$. *$p < 0.05$ and #$p < 0.05$ (n = 6). The data are represented as the mean ± SEM values.

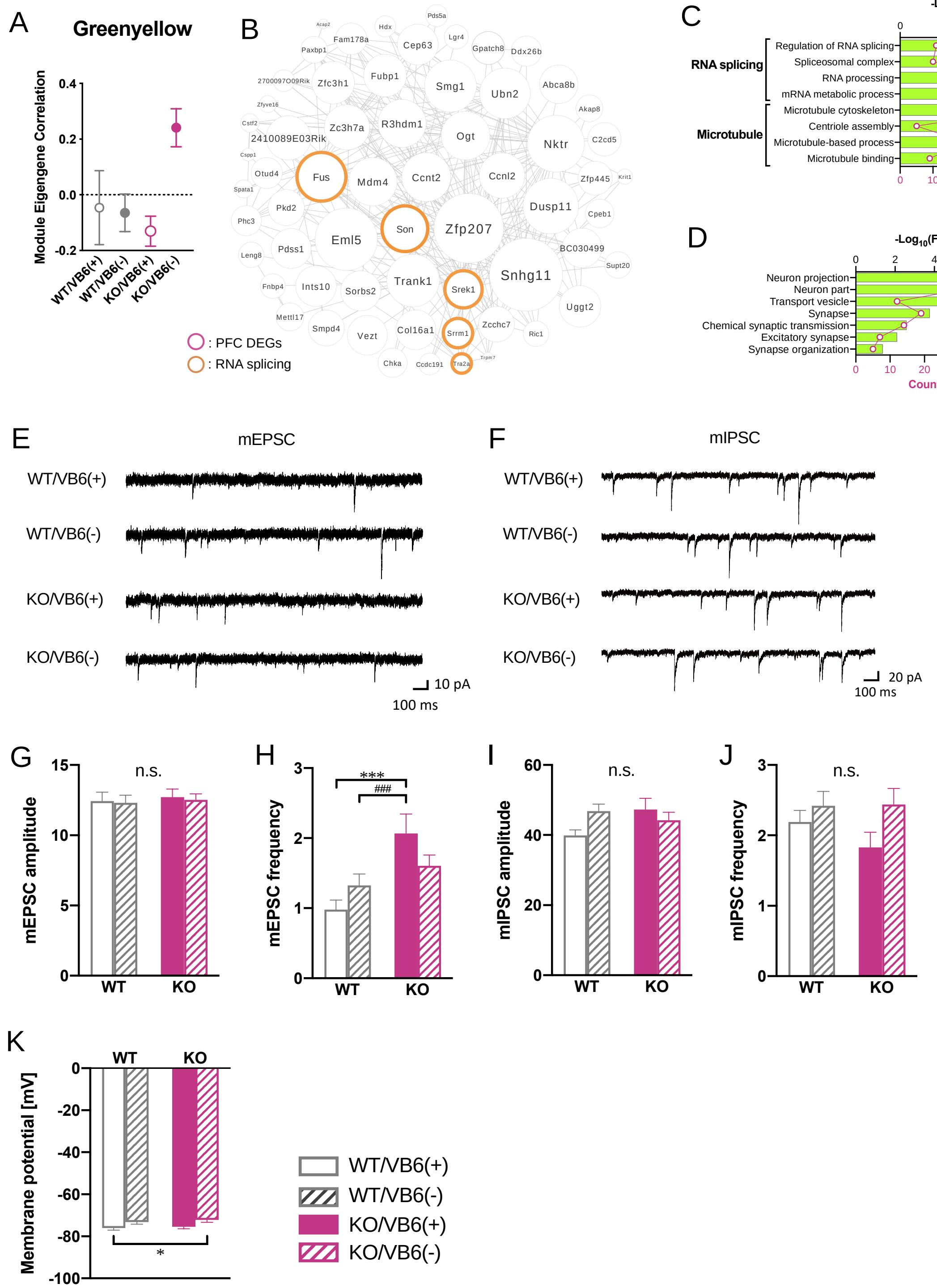
A
Greenyellow
Module Eigengene Correlation
WT/VB6(+)
WT/VB6(-)
KO/VB6(+)
KO/VB6(-)
B
: PFC DEGs
: RNA splicing
C
-Log10(FDR)
RNA splicing
Regulation of RNA splicing
Spliceosomal complex
RNA processing
mRNA metabolic process
Microtubule
Microtubule cytoskeleton
Centriole assembly
Microtubule-based process
Microtubule binding
Count
D
-Log10(FDR)
Neuron projection
Neuron part
Transport vesicle
Synapse
Chemical synaptic transmission
Excitatory synapse
Synapse organization
Count
E
mEPSC
10 pA
100 ms
F
mIPSC
20 pA
100 ms
G
mEPSC amplitude
n.s.
WT
KO
H
mEPSC frequency
***
###
I
mIPSC amplitude
J
mIPSC frequency
K
Membrane potential [mV]
*

***Supplementary Figure 4: Weighted gene coexpression network analysis (WGCNA) and electrophysiological changes in the PFC***

(A) Module eigengene of greenyellow module significantly associated with KO/VB6(-) group. (B) Visualization of the top 250 connections ranked by weighted topological overlap values for the greenyellow module. Node size corresponds to the number of edges (degree). GO analyses of genes in (C) the greenyellow and (D) genes indicated differential exon usage in the PFC of KO/VB6(-) are shown. Representative data showing the effect of *Glo1* deletion and VB6-deficiency on (E) mEPSC and (F) mIPSC in the PFC are shown. Amplitude of (G) mEPSC and (I) mIPSC, and frequency of (H) mEPSC and (J) mIPSC are shown. (K) Membrane potential were measured. Two-way ANOVA: (G) $F_{\text{Interaction}(1, 111)} = 0.01$, $p > 0.05$; $F_{\text{Genotype}(1, 111)} = 0.20$, $p > 0.05$; $F_{\text{VB6}(1, 111)} = 0.07$, $p > 0.05$, (H) $F_{\text{Interaction}(1, 109)} = 4.08$, $p < 0.05$; $F_{\text{Genotype}(1, 109)} = 11.7$, $p < 0.001$; $F_{\text{VB6}(1, 109)} = 0.09$, $p > 0.05$, (I) $F_{\text{Interaction}(1, 127)} = 4.56$, $p < 0.05$; $F_{\text{Genotype}(1, 127)} = 1.06$, $p > 0.05$; $F_{\text{VB6}(1, 127)} = 0.69$, $p > 0.05$, (J) $F_{\text{Interaction}(1, 124)} = 0.84$, $p > 0.05$; $F_{\text{Genotype}(1, 124)} = 0.70$, $p > 0.05$; $F_{\text{VB6}(1, 124)} = 4.15$, $p < 0.05$, (K) $F_{\text{Interaction}(1, 93)} = 0.03$, $p > 0.05$; $F_{\text{Genotype}(1, 93)} = 0.66$, $p > 0.05$; $F_{\text{VB6}(1, 93)} = 9.28$, $p < 0.01$. *$p < 0.05$, ***$p < 0.001$, and ###$p < 0.001$. The data are represented as the mean ± SEM values.
