## Supplemental Methods for "Impairment of methylglyoxal detoxification systems causes mitochondrial dysfunction and schizophrenia-like behavioral deficits"

**Supplementary Methods**

***Open field test***

The open-field was a square with grey walls (W50 × D50 × H40 cm), placed in a dark, sound-attenuated room. Each mouse was placed in one corner of the open field, and it was allowed to explore the environment freely for 10 min. Time in center (16.7 x 16.7 cm) and traveled distance were determined using the EthoVision system (Neuroscience Idea Co. Ltd., Osaka, Japan).

***Y-maze test***

Short-term memory was assessed by recording spontaneous alternation behavior during a single session in a Y-maze. Each arm was 40 cm long, 12 cm high, 3 cm wide at the bottom, 10 cm wide at the top, and converged in an equilateral triangular central area. Each mouse, naïve to the maze, was placed at the end of one arm and allowed to move freely through the maze during a 10 min session, and the number of arm entries was counted. Each series of arm entries was recorded visually, and an arm entry was defined as when the hind paws of the mouse were completely within the arm. Alternation was defined as successive entries into the three arms on the overlapping triplet sets. The percentage alternation was calculated using the following formula: (number of alternations)/(total number of arm entries-2) × 100 (%).

***Differential exon usage analysis***

Differential expression between KO/VB6(-) and WT/VB6(+) samples was assessed using DEXseq package in R[1] based on mm10 Gencode (version vM11) exonic annotation file gtf file. We applied the following method on all the pairwise comparison between WT and KO, with/without Vitamin in the diet.

Using SVA package in R[2], we identified potential unwanted surrogate variables (2 surrogate variables).

Linear regression was performed using DEXseq to remove covariate variables:

$$Exon\ expression \sim SVA1 + SVA2 + Batch$$

We retained an exon differentially expressed for an FDR <= 0.05.

***Electrophysiology***

Under $CO_2$ anesthesia, coronal prefrontal slices of 300 μm thickness were prepared with a vibratome slicer (VT-1200S, Leica) from the adult mice using the following cutting solution; 120 mM Choline Cl, 28 mM $NaHCO_3$, 1.25 mM $NaH_2PO_4$, 2 mM KCl, 25 mM glucose, 1 mM $CaCl_2$, 8 mM $MgCl_2$, bubbled with 95% $O_2$ and 5% $CO_2$ (Narushima et al., 2006) . The slices were incubated for 30-60 min at room temperature with artificial cerebrospinal fluid (ACSF) containing 125 mM NaCl, 2.5 mM KCl, 2 mM $CaCl_2$, 1 mM $MgSO_4$, 1.25 mM $NaH_2PO_4$, 26 mM $NaHCO_3$, and 20 mM glucose bubbled with 95% $O_2$ and 5% $CO_2$ (Uesaka et al., 2014).

Whole-cell patch clamp recording was performed in a blind manner to experimental groups. We identified pyramidal neurons of layer 2/3 in the medial prefrontal cortex (mPFC) morphologically with Olympus BX51WI microscope (Olympus). The pipet resistance was 2.4-5 MΩ when filled with an internal solution containing 130 mM potassium D-gluconate, 6 mM KCl, 10 mM NaCl, 10 mM HEPES, 0.16 mM $CaCl_2$, 2 mM $MgCl_2$, 0.5 mM EGTA, 4 mM Na-ATP, and 0.4 mM Na-GTP (pH 7.3, adjusted with KOH) (Maejima et al., 2005) for recording miniature excitatory postsynaptic currents (mEPSCs) and for current-clamp recordings. For recording miniature inhibitory postsynaptic currents (mIPSCs), an internal solution with the following composition was used: 145 mM KCl, 10 mM HEPES, 10 mM EGTA, 0.16 mM $CaCl_2$, 2 mM $MgCl_2$, 5 mM Mg-ATP, and 0.2 mM Na-GTP (pH 7.2, adjusted with KOH) (Ade et al., 2008). All the data were recorded at 30-32°C with an EPC-10 amplifier (HEKA Elektronik) and filtered at 2.9 kHz and digitized at 20 kHz. The access resistance was compensated by 70% and he liquid junction potential was corrected. mEPSCs and mIPSCs were recorded at the holding potential of −70 mV in the presence of 0.5 μM tetrodotoxin (TTX) (Nacalai Tesque) together with 0.1 mM picrotoxin (PTX) (for mEPSCs) or 10 μM NBQX and 50 μM D-AP5 (for mIPSCs). Mini Analysis program (Synaptosoft) was used for analyzing mEPSCs and mIPSCs. Individual synaptic events were detected by semi-automatic method or eye when the amplitude was larger than 5 pA and the rise time was faster than 3 ms in a blind manner, and after finishing all the analysis genotypes and conditions were assigned to experimental groups.
